# Gene duplication and retrotransposition diversify the antiviral repertoire of macaque IFITM proteins

**DOI:** 10.64898/2026.06.11.731520

**Authors:** Deepthi Kappala, Alexis Sauer, Emma Atwood, Yuexiu Zhang, Anuja Pimplapure, Naveen Bektas-Jolly, Lohitha Gujjari, Kasturi Chandra, Yogesh Saini, Jacob S. Yount, Amol Suryawanshi, Jianrong Li, Amit Sharma

**Author notes:** Address correspondence to: Amit Sharma, PhD Department of Population Health and Pathobiology, North Carolina State University, Raleigh, NC 27607.

## Abstract

Interferon-induced transmembrane (IFITM) proteins are broad-spectrum antiviral restriction factors that inhibit viral entry of diverse enveloped viruses. Comparative genomic studies have revealed extensive lineage-specific diversification of IFITM genes, yet the functional consequences of this diversification remain poorly understood. We previously identified an expanded IFITM repertoire in macaques consisting of the canonical IFITM proteins, IFITM1 and IFITM3, a duplicated IFITM3 paralog (IFITM3A), and two retrotransposed IFITM3-derived genes (IFITM3-R1 and IFITM3-R2). Here, we thoroughly characterized the antiviral activities, intracellular localization, and mechanisms of regulation of canonical and non-canonical macaque IFITMs against vesicular stomatitis virus (VSV), influenza A virus (IAV), Sendai virus, and HIV-1. IFITM3A exhibited enhanced antiviral activity relative to IFITM3, particularly against VSV and HIV-1. Comparative mutational analyses identified amino acid substitutions in IFITM3 that contribute to the enhanced antiviral phenotype of IFITM3A. In contrast, the retrocopy IFITM3-R1 exhibited markedly reduced expression due to lysosome-dependent protein turnover mediated by a PPxY motif and a unique lysine residue (K51). Alteration of these determinants increased IFITM3-R1 expression and selectively enhanced restriction of VSV and IAV. Although several macaque IFITMs reduced HIV-1 infectivity when expressed in producer cells, none significantly inhibited HIV-1 infection in target cells. Furthermore, differential incorporation of IFITMs into HIV-1 virions did not consistently correlate with antiviral activity, indicating that virion incorporation alone is insufficient to explain HIV-1 restriction. Together, these findings demonstrate that gene duplication and retrotransposition have generated a functionally diverse IFITM repertoire in macaques and provide insight into how evolutionary diversification expands innate antiviral defenses in primates.

**Importance:** IFITM proteins are broad-spectrum antiviral restriction factors that inhibit infection of numerous enveloped viruses. Although IFITM genes have undergone extensive diversification during mammalian evolution, the functional consequences of this diversification remain poorly understood. Here, we show that expansion of the macaque IFITM locus through gene duplication and retrotransposition generated proteins with distinct antiviral activities, intracellular localization patterns, and regulatory mechanisms. We identify amino acid determinants that contribute to the enhanced antiviral activity of the duplicated paralog IFITM3A and demonstrate that lysosome-dependent turnover mediated by a PPxY motif and a unique lysine residue limits expression and antiviral activity of the retrocopy IFITM3-R1. We further show that macaque IFITMs inhibit HIV-1 predominantly through producer-cell-dependent mechanisms and reveal that IFITM virion incorporation alone does not predict antiviral potency. These findings provide mechanistic insight into how restriction factor diversification expands innate antiviral defenses and shapes host-virus interactions in primates.

## Introduction

Interferon-induced transmembrane (IFITM) proteins are among the first lines of intrinsic antiviral defense and act by restricting viral entry at cellular and endosomal membranes (1). Since their identification as broad-spectrum restriction factors, IFITMs have been shown to inhibit numerous enveloped viruses, including influenza A virus (IAV), HIV-1, dengue virus, Ebola virus, and various coronaviruses (2–5). By targeting early stages of membrane fusion, IFITMs impose a potent barrier to infection that can influence viral tropism, pathogenicity, and host range. Because viral entry is a major determinant of zoonotic transmission, IFITMs are also increasingly recognized as important host determinants shaping cross-species transmission and viral emergence (6–8).

The continuous conflict between viral pathogens and host restriction factors drives a dynamic evolutionary “arms race”, in which viruses evolve mechanisms to evade restriction while hosts diversify antiviral defenses to maintain protection (9). Gene duplication represents one of the most effective evolutionary strategies for expanding antiviral repertoires. Duplicated restriction factors can acquire novel antiviral specificities, altered subcellular localization, or enhanced potency while preserving essential ancestral functions (10). Such diversification enables hosts to simultaneously target multiple viruses or constrain viral escape through mechanistically distinct antiviral activities. In primates, this evolutionary principle has shaped several innate immune gene families, including APOBEC3, Mx GTPases, TRIM5, and IFITMs, highlighting the importance of gene expansion in antiviral adaptation (10–15).

Among primates, macaques represent a particularly informative model for investigating IFITM evolution and function because they encode a substantially expanded and diversified IFITM locus compared with humans (15–17). In our previous evolutionary and genomic analysis of macaque IFITM genes, we identified extensive lineage-specific duplications and retrotransposition events that have reshaped the macaque IFITM repertoire (16). In contrast to humans, which encode the canonical IFITM1, IFITM2, and IFITM3 genes, macaques lack IFITM2 but instead encode a duplicated IFITM3 paralog, IFITM3A. IFITM3A retains an intact open reading frame comparable in length to IFITM3 yet diverges substantially at the amino acid level, suggesting functional specialization following duplication. In addition, macaque genomes harbor multiple non-canonical IFITM retrocopies, including IFITM3-R1 and IFITM3-R2, as well as several pseudogenes, indicating recurrent evolutionary innovation within this locus. These findings raised the possibility that macaques have evolved a broader and potentially more specialized IFITM antiviral network than humans.

Despite the remarkable diversification of macaque IFITM genes, the antiviral activities, regulatory mechanisms, and functional consequences of this expanded repertoire remain poorly understood. In particular, it is unknown whether duplicated and retrotransposed IFITMs have retained antiviral activity, evolved virus-specific functions, or acquired distinct mechanisms governing intracellular trafficking, protein stability, and antiviral restriction. Understanding how these genes contribute to antiviral defense is important not only for defining the evolutionary consequences of IFITM expansion, but also for understanding innate immune barriers that influence susceptibility to viral infection in macaques, which are widely used as models for HIV/AIDS, influenza, and emerging viral diseases.

Here, we systematically characterize the antiviral repertoire of canonical and non-canonical macaque IFITM proteins against a diverse panel of enveloped RNA viruses, including vesicular stomatitis virus (VSV), IAV, Sendai virus (SeV), and HIV-1. We demonstrate that macaque IFITMs exhibit distinct antiviral specificities that are associated with differences in intracellular localization, protein stability, and protein degradation pathway. We identify amino acid determinants that contribute to the enhanced antiviral activity of the duplicated paralog IFITM3A and uncover mechanisms that limit the expression and antiviral potential of the retrocopied IFITM3-R1 protein. In addition, we demonstrate that macaque IFITMs inhibit HIV-1 infectivity through producer-cell-dependent mechanisms and reveal that virion incorporation alone does not predict antiviral activity. Together, these findings define the functional consequences of IFITM diversification in macaques and provide insight into how gene duplication and retrotransposition expand innate antiviral defenses in non-human primates.

## Results

### Canonical and non-canonical macaque IFITMs exhibit distinct antiviral activities and intracellular localization patterns

We previously identified an expanded repertoire of IFITM genes in macaques consisting of the canonical IFITM proteins IFITM1 and IFITM3, a duplicated IFITM3 paralog (IFITM3A), and two retrotransposed IFITM3-derived genes (IFITM3-R1 and IFITM3-R2) located outside the canonical IFITM locus (16) (Figs. S1 and S2). To investigate the functional consequences of this diversification, we compared the antiviral activities of canonical and non-canonical macaque IFITMs against a panel of enveloped RNA viruses.

As a reference, we first characterized human IFITM1, IFITM2, and IFITM3. All three proteins were readily detected by immunoblotting and expressed at comparable levels following transient expression in HEK293T cells (Fig. 1A). Next, we determined the antiviral activities of human IFITMs against VSV, IAV, SeV, and HIV-1. Consistent with previous studies (18–22), human IFITMs restricted VSV and IAV infections by ∼20–80%, whereas SeV remained largely resistant (Fig. 1B–D). In contrast, expression of human IFITMs in target cells had no observable effect on HIV-1 infection (Fig. 1E), indicating that HIV-1 entry is relatively insensitive to IFITM-mediated restriction. However, expression of IFITM2 and IFITM3 in HIV-1 producer cells reduced virion infectivity by ∼60–70%, whereas IFITM1 had no observable effect on virion infectivity (Fig. 1F). Consistent with a producer-cell mechanism of restriction (23–26), all three human IFITMs were incorporated into HIV-1 particles (Fig. 1G). Confocal microscopy revealed localization patterns consistent with previous reports, with IFITM1 predominantly enriched at the plasma membrane and IFITM2 and IFITM3 predominantly localized within endosomal compartments (Fig. 1H–J).

**Figure 1.**
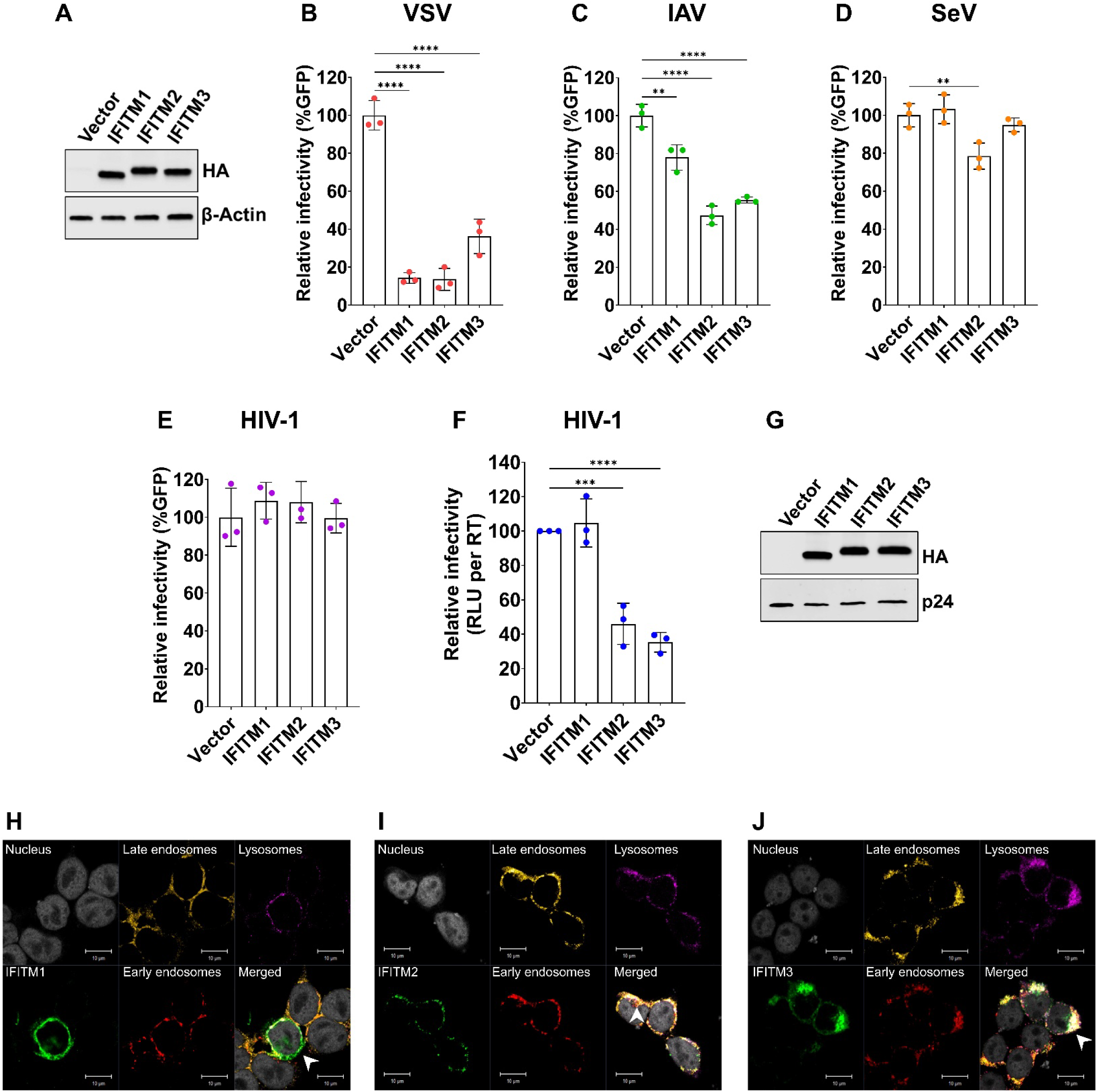
Human IFITM proteins display distinct antiviral activities and subcellular localization patterns. **(A)** Expression of HA-tagged human IFITM1, IFITM2, and IFITM3 in HEK293T cells. Western blot analysis of HEK293T cells expressing the indicated HA-tagged IFITM proteins or an empty vector control (Vector) using anti-HA and anti-β-actin antibodies. **(B–E)** Effect of human IFITMs on viral infectivity. HEK293T cells expressing the indicated IFITM proteins were infected with GFP-expressing vesicular stomatitis virus (VSV) **(B)**, influenza A virus (IAV) **(C)**, Sendai virus (SeV) **(D)**, or pseudotyped HIV-1 **(E)**. Infection was quantified by flow cytometry as the percentage of GFP-positive cells and normalized to the empty vector control. **(F)** Effect of human IFITMs on HIV-1 infectivity. HIV-1 virions were produced in HEK293T cells expressing the indicated IFITM proteins. Infectivity was determined on TZM-bl target cells by measuring relative light units (RLU) normalized to reverse transcriptase (RT) activity and expressed relative to the empty vector control. **(G)** Incorporation of human IFITMs into HIV-1 virions. Virions produced in the presence of the indicated IFITM proteins were analyzed by Western blot using anti-HA and anti-p24 antibodies. **(H–J)** Subcellular localization of human IFITM1 **(H)**, IFITM2 **(I)**, and IFITM3 **(J)**. HEK293T cells expressing the indicated proteins were stained for nuclei, IFITM (HA), early endosomes (EEA1), late endosomes (Rab7), and lysosomes (LAMP-1). Representative confocal images are shown. Arrowheads indicate regions of IFITM localization. Scale bars, 10 μm. For panels B–F, bars represent means from three independent experiments, with individual data points shown. Error bars indicate standard deviations. Statistical significance was determined by one-way ANOVA with multiple-comparison correction. *, P < 0.05; **, P < 0.01; ***, P < 0.001; ****, P < 0.0001.

We next examined the antiviral activities of macaque IFITMs. Immunoblot analysis confirmed robust and comparable expression of IFITM1, IFITM3, IFITM3A, and IFITM3-R2 in HEK293T cells, whereas IFITM3-R1 was consistently expressed at substantially lower levels (Fig. 2A). Functional analysis revealed marked differences among macaque IFITMs. IFITM3A displayed the strongest antiviral activity, reducing VSV infection by ∼70%, whereas both IFITM3 and IFITM3A reduced IAV infection by ∼30% (Fig. 2B–C). Similar to human IFITMs, SeV remained largely resistant to inhibition by all macaque IFITMs (Fig. 2D). Notably, expression of macaque IFITMs in target cells produced no inhibition of HIV-1 infection (Fig. 2E), mirroring the phenotype observed with human IFITMs and suggesting that macaque IFITMs do not efficiently block HIV-1 entry. In contrast, expression of IFITM3A and IFITM3-R2 in HIV-1 producer cells reduced virion infectivity by ∼30% (Fig. 2F–G), supporting a mechanism in which macaque IFITMs influence HIV-1 infectivity through effects on progeny virions rather than target-cell entry.

**Figure 2.**
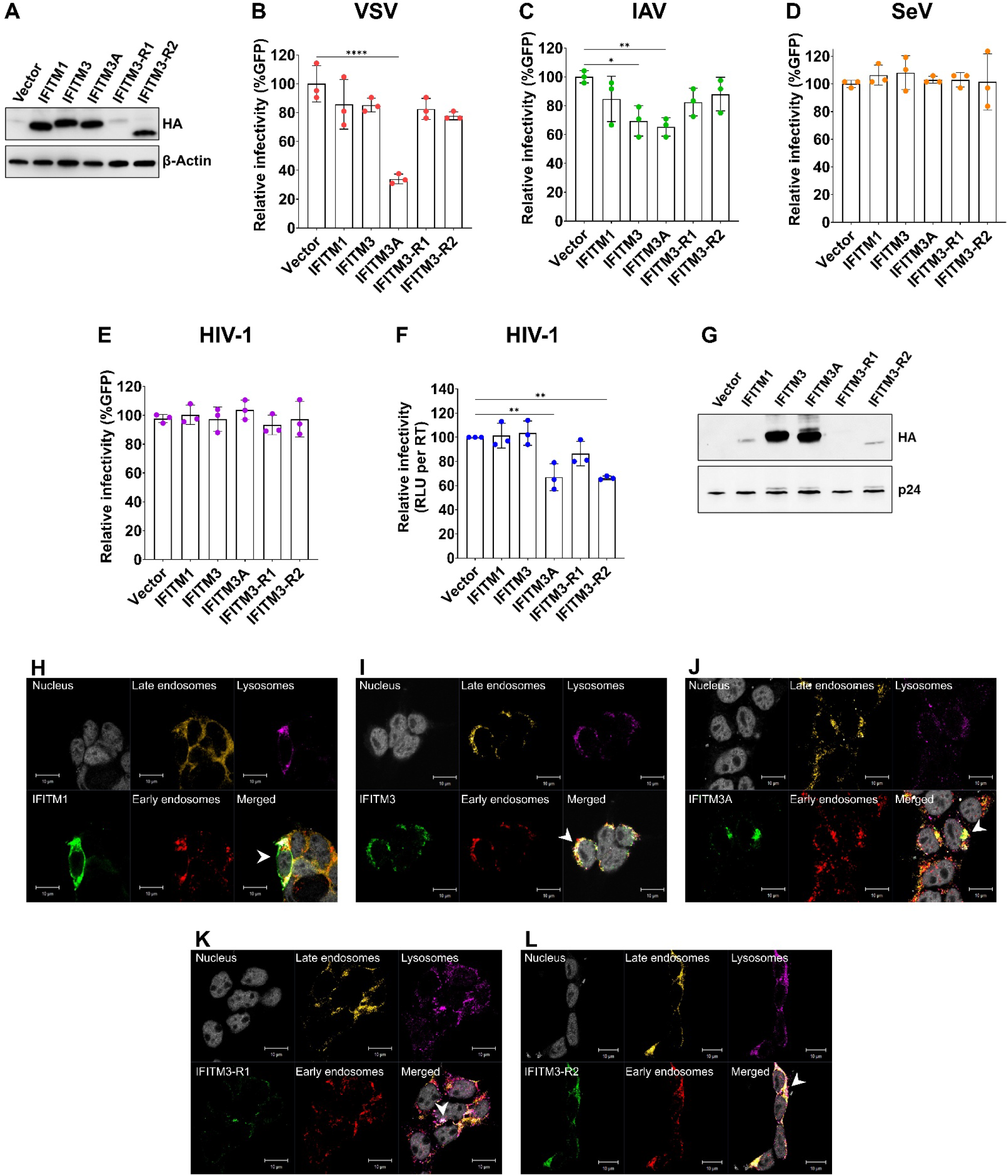
Canonical and non-canonical macaque IFITM proteins differ in antiviral activity and intracellular localization. **(A)** Expression of HA-tagged macaque IFITM1, IFITM3, IFITM3A, IFITM3-R1, and IFITM3-R2 in HEK293T cells. Western blot analysis of cells expressing the indicated IFITM proteins or an empty vector control (Vector) using anti-HA and anti-β-actin antibodies. **(B–E)** Effect of macaque IFITMs on viral infectivity. HEK293T cells expressing the indicated IFITM proteins were infected with GFP-expressing VSV **(B)**, IAV **(C)**, SeV **(D)**, or pseudotyped HIV-1 **(E)**. Infection was quantified by flow cytometry as the percentage of GFP-positive cells and normalized to the empty vector control. **(F)** Effect of macaque IFITMs on HIV-1 infectivity. HIV-1 virions were produced in HEK293T cells expressing the indicated IFITM proteins. Infectivity was determined on TZM-bl target cells by measuring RLU normalized to RT activity and expressed relative to the empty vector control. **(G)** Incorporation of macaque IFITMs into HIV-1 virions. Virions produced in the presence of the indicated IFITM proteins were analyzed by Western blot using anti-HA and anti-p24 antibodies. **(H–L)** Subcellular localization of macaque IFITM1 **(H)**, IFITM3 **(I)**, IFITM3A **(J)**, IFITM3-R1 **(K)**, and IFITM3-R2 **(L)**. HEK293T cells expressing the indicated proteins were stained for nuclei, IFITM (HA), early endosomes (EEA1), late endosomes (Rab7), and lysosomes (LAMP-1). Representative confocal images are shown. Arrowheads indicate regions of IFITM localization. Scale bars, 10 μm. For panels B–F, bars represent means from three independent experiments, with individual data points shown. Error bars indicate standard deviations. Statistical significance was determined by one-way ANOVA with multiple-comparison correction. *, P < 0.05; **, P < 0.01; ***, P < 0.001; ****, P < 0.0001.

To determine whether differences in antiviral activity correlated with altered intracellular localization and trafficking, we examined the subcellular localization of macaque IFITMs by confocal microscopy. IFITM1 localized predominantly to the plasma membrane, whereas IFITM3 accumulated primarily within Rab7-positive late endosomes (Fig. 2H–I). In contrast, IFITM3A displayed a broader endosomal distribution, localizing to both EEA1-positive early endosomes and Rab7-positive late endosomes (Fig. 2J). IFITM3-R2 exhibited a diffuse distribution, while expression of IFITM3-R1 was insufficient for reliable imaging (Fig. 2K–L). Together, these findings demonstrate that expansion of the macaque IFITM repertoire has generated proteins with distinct antiviral activities and intracellular localization patterns.

### Amino acid substitutions in IFITM3 contribute to the enhanced antiviral activity of IFITM3A

Because IFITM3A exhibited enhanced antiviral activity relative to IFITM3, particularly against VSV (Fig. 2B), we sought to identify amino acid determinants that contribute to this phenotype. Sequence comparison of macaque IFITM3A, macaque IFITM3, and human IFITM2 revealed several non-conservative substitutions at positions previously implicated in IFITM trafficking and antiviral activity (Fig. S3). Based on these analyses, we selected three candidate residues for functional evaluation.

The first substitution, F8V, is located within the highly conserved N-terminal region of IFITM proteins. Human IFITM3 and macaque IFITM3 contain a conserved double-phenylalanine motif at amino acid positions 8 and 9, whereas IFITM3A encodes a valine at position 8 and retains a single phenylalanine at position 9. Notably, the double-phenylalanine motif is a conserved feature of IFITM3 proteins across multiple mammalian species (27), suggesting an important functional role. The presence of a divergent residue at this position in IFITM3A suggested that this modification of the N-terminus may contribute to its distinct antiviral activity.

The second substitution, P70T, is located within the intramembrane domain, a region critical for IFITM3-mediated restriction of viral membrane fusion (28). Whereas human IFITM3 and macaque IFITM3 encode a proline at amino acid position 70, both human IFITM2 and macaque IFITM3A encode a threonine at this position. Because IFITM2 exhibited modest antiviral activity against SeV compared with IFITM3 (Fig. 1D), we hypothesized that substitution of proline 70 may contribute to the altered antiviral phenotype of IFITM3A.

The third substitution, I108V, is located within the conserved C-terminal region. This residue was selected based on a previous study demonstrating that mutations within the neighboring 103–108 region influence IFITM3 stability and virion incorporation (29). Specifically, alterations within this region were reported to enhance incorporation of human IFITM3 into HIV-1 virions despite reducing protein stability. Because IFITM3A naturally encodes a valine at position 108, we reasoned that this substitution could contribute to differences in anti-HIV activity between IFITM3 and IFITM3A.

To determine whether these naturally occurring amino acid differences contribute to the enhanced antiviral activity of IFITM3A, each substitution was individually introduced into macaque IFITM3, and the resulting mutants were evaluated for antiviral activity, intracellular localization, and incorporation into HIV-1 virions. Immunoblot analysis confirmed robust and comparable expression of IFITM3, IFITM3A, and the three IFITM3 point mutants in HEK293T cells (Fig. 3A). Both F8V and P70T enhanced the ability of IFITM3 to restrict VSV infection, reducing infectivity to levels comparable to those observed with IFITM3A (Fig. 3B). Although these mutants also reduced IAV infection, their effects were not substantially greater than those observed with wild-type IFITM3, which already restricted IAV efficiently (Fig. 3C). Neither mutation appreciably altered HIV-1 infectivity when expressed in producer cells (Fig. 3D). In contrast, the I108V mutation selectively reduced HIV-1 infectivity by ∼30%, phenocopying the antiviral activity of IFITM3A (Fig. 3D).

**Figure 3.**
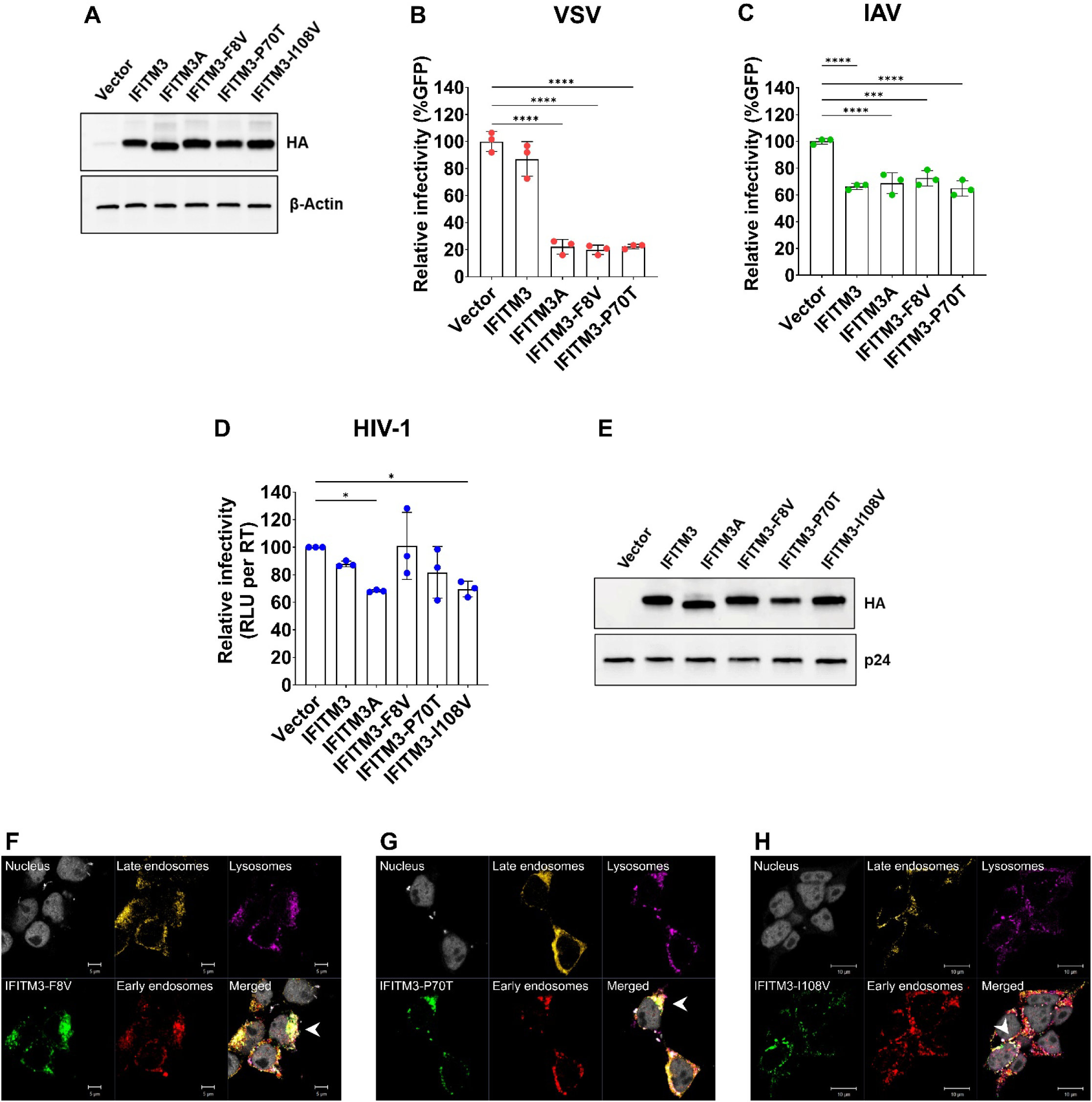
Determinants of antiviral activity of macaque IFITM3 proteins. **(A)** Expression of HA-tagged macaque IFITM3, IFITM3A, and the indicated IFITM3 variants (F8V, P70T, and I108V) in HEK293T cells. Western blot analysis of cells expressing the indicated proteins or an empty vector control (Vector) using anti-HA and anti-β-actin antibodies. **(B–C)** Effect of IFITM3 variants on viral infectivity. HEK293T cells expressing the indicated IFITM3 proteins were infected with GFP-expressing VSV **(B)** or IAV **(C)**. Infection was quantified by flow cytometry as the percentage of GFP-positive cells and normalized to the empty vector control. **(D)** Effect of IFITM3 variants on HIV-1 infectivity. HIV-1 virions were produced in HEK293T cells expressing the indicated IFITM3 proteins. Infectivity was determined on TZM-bl target cells by measuring RLU normalized to RT activity and expressed relative to the empty vector control. **(E)** Incorporation of IFITM3 variants into HIV-1 virions. Virions produced in the presence of the indicated IFITM3 proteins were analyzed by Western blot using anti-HA and anti-p24 antibodies. **(F–H)** Subcellular localization of IFITM3-F8V **(F)**, IFITM3-P70T **(G)**, and IFITM3-I108V **(H)**. HEK293T cells expressing the indicated proteins were stained for nuclei, IFITM3 (HA), early endosomes (EEA1), late endosomes (Rab7), and lysosomes (LAMP-1). Representative confocal images are shown. Arrowheads indicate regions of IFITM localization. Scale bars, 10 μm. For panels B–D, bars represent means from three independent experiments, with individual data points shown. Error bars indicate standard deviations. Statistical significance was determined by one-way ANOVA with multiple-comparison correction. *, P < 0.05; **, P < 0.01; ***, P < 0.001; ****, P < 0.0001.

To investigate potential mechanism(s) underlying these virus-specific effects, we examined virion incorporation and intracellular localization of the IFITM3 mutants. All three variants remained detectable in purified HIV-1 virions (Fig. 3E) and displayed intracellular localization patterns resembling IFITM3A, with partial overlap with the early and late endosomal compartments (Fig. 3F–H). These findings indicate that specific amino acid residues contribute to restriction of different viruses and collectively account for enhanced antiviral phenotypes of IFITM3A.

### A PPxY motif limits expression of non-canonical IFITM3 retrocopies

Among the non-canonical IFITMs, IFITM3-R1 consistently exhibited markedly reduced protein expression despite repeated optimization of transfection conditions (Fig. 2A). To identify the determinant(s) responsible for this phenotype, we compared the amino acid sequences of canonical and non-canonical macaque IFITMs and identified a conserved N-terminal PPxY motif within IFITM3-R1 and IFITM3-R2 (Fig. 4A).

**Figure 4.**
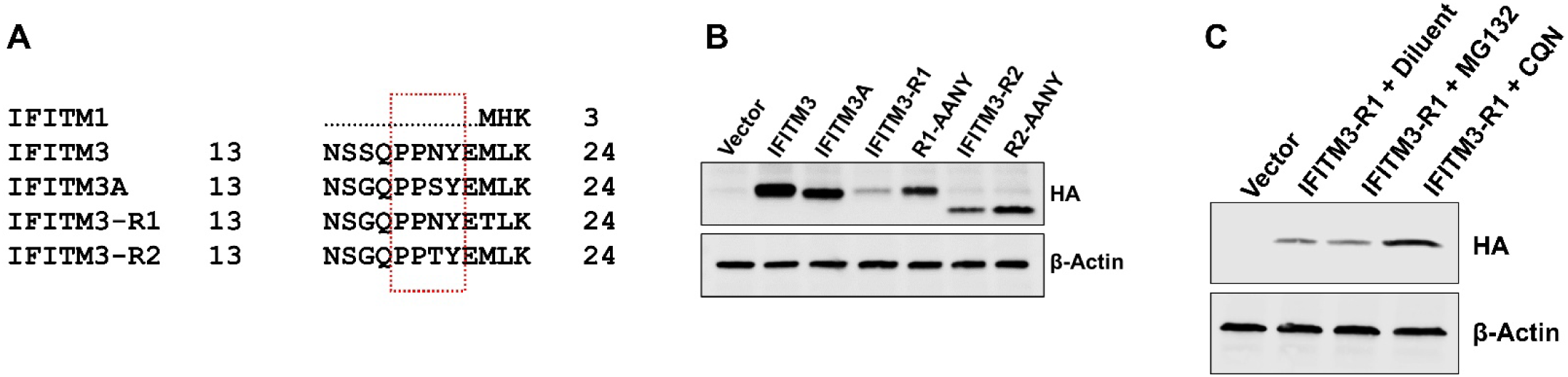
A PPxY motif limits expression of non-canonical macaque IFITM3 proteins. **(A)** Alignment of the N-terminal sequences of macaque IFITM proteins. PPxY motifs that differ between canonical and non-canonical IFITM proteins are highlighted. **(B)** Mutation of the PPxY motif increases expression of IFITM3-R1 and IFITM3-R2. HEK293T cells expressing HA-tagged IFITM3, IFITM3A, IFITM3-R1, IFITM3-R2, or the indicated AANY-substitution mutants were analyzed by Western blot using anti-HA and anti-β-actin antibodies. **(C)** Chloroquine treatment restores IFITM3-R1 protein expression. HEK293T cells expressing HA-tagged IFITM3-R1 were treated with vehicle control, MG132 (10 μM), or chloroquine (CQN; 10 μM) for 24 hours. Cell lysates were analyzed by Western blot using anti-HA and anti-β-actin antibodies.

To determine whether the PPxY motif contributes to reduced retrocopy expression, we mutated the PPxY motif to AANY in both IFITM3-R1 and IFITM3-R2. Disruption of the PPxY motif increased expression of both retrocopies, particularly IFITM3-R1 (Fig. 4B), indicating that the PPxY motif negatively regulates protein accumulation.

Because PPxY motifs frequently mediate interactions with cellular trafficking and protein degradation machinery (30, 31), we next investigated whether reduced IFITM3-R1 expression reflected enhanced protein turnover. We employed pharmacological inhibitors to block the autophagy-lysosome pathway and the ubiquitin-proteasome system in HEK293T cells expressing IFITM3-R1. Treatment of IFITM3-R1-expressing cells with the lysosomal inhibitor chloroquine markedly increased IFITM3-R1 protein levels, whereas the proteasome inhibitor MG132 had no effect on protein levels (Fig. 4C). These findings indicate that IFITM3-R1 is preferentially targeted for lysosomal degradation and suggest that the PPxY motif contributes to this process.

### Disruption of the PPxY motif selectively enhances antiviral activity of IFITM3 retrocopies

We next examined whether increased IFITM3 retrocopy expression resulting from disruption of the PPxY motif translated into enhanced antiviral activity. Mutation of the PPxY motif selectively increased the antiviral activity of both retrocopies, although the magnitude and specificity of these effects varied among viruses.

For IFITM3-R1, disruption of the PPxY motif significantly enhanced inhibition of IAV infection, restoring antiviral activity to levels comparable to those observed for canonical IFITM3 proteins (Fig. 5B). In contrast, the IFITM3-R2 AANY mutant exhibited enhanced restriction of VSV infection, although its activity remained lower than that of IFITM3A (Fig. 5A). Interestingly, the IFITM3-R2 mutant also acquired measurable activity against SeV, a virus largely resistant to inhibition by the other IFITMs examined in this study (Fig. 5C).

**Figure 5.**
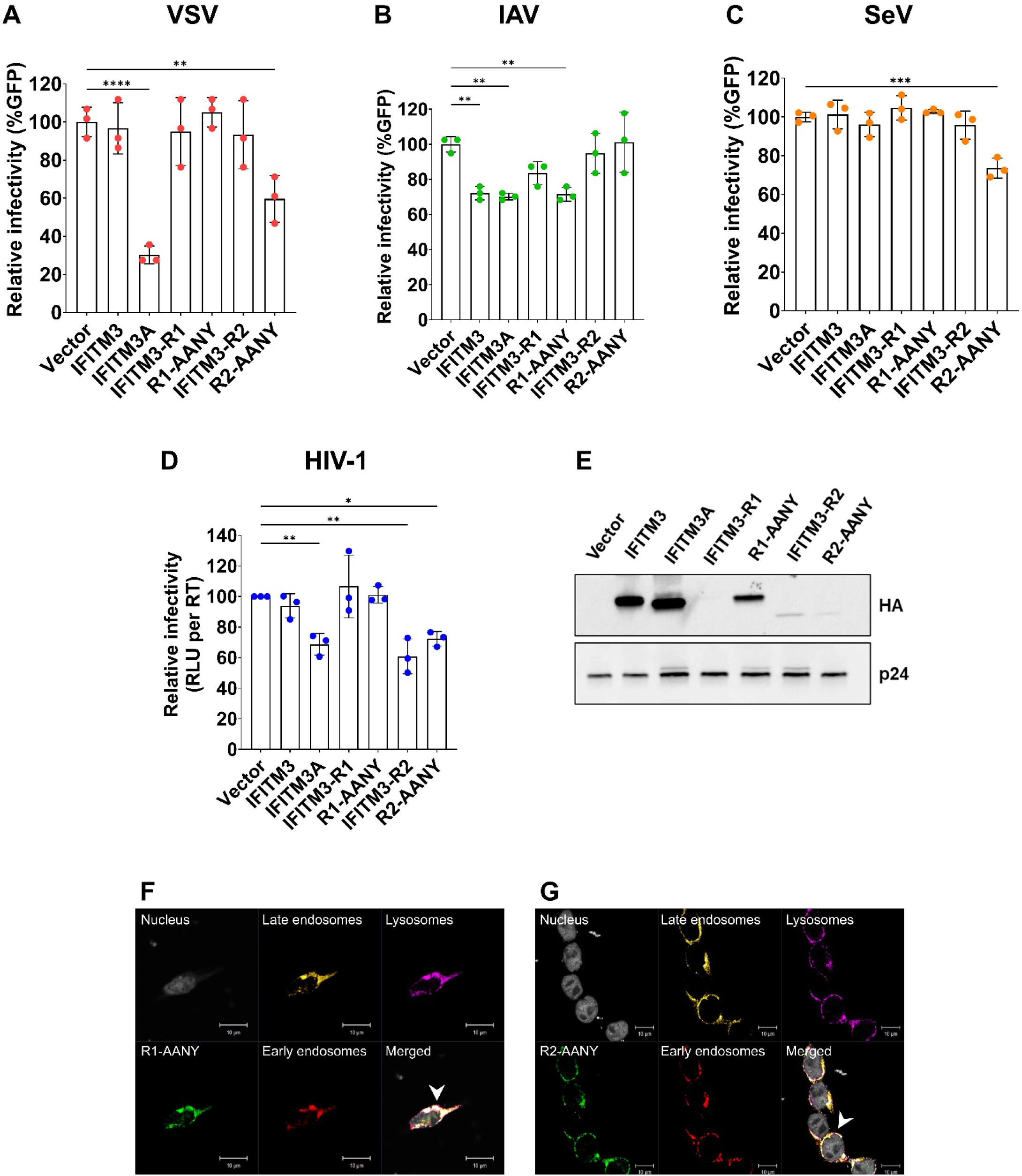
Disruption of the PPxY motif enhances antiviral activity of non-canonical macaque IFITM3 proteins. **(A–C)** Effect of PPxY motif mutations on antiviral activity. HEK293T cells expressing the indicated canonical and non-canonical IFITM3 proteins, the corresponding AANY-substitution mutants, or an empty vector control were infected with GFP-expressing VSV **(A)**, IAV **(B)**, or SeV **(C)**. Infection was quantified by flow cytometry as the percentage of GFP-positive cells and normalized to the empty vector control. **(D)** Effect of PPxY motif mutations on HIV-1 infectivity. HIV-1 virions were produced in HEK293T cells expressing the indicated IFITM proteins or mutants. Infectivity was determined on TZM-bl target cells by measuring RLU normalized to RT activity and expressed relative to the empty vector control. **(E)** Incorporation of IFITM3 variants and corresponding AANY-substitution mutants into HIV-1 virions. Virions produced in the presence of the indicated IFITM proteins were analyzed by Western blot using anti-HA and anti-p24 antibodies. **(F–G)** Subcellular localization of IFITM3-R1-AANY **(F)** and IFITM3-R2-AANY **(G)**. HEK293T cells expressing the indicated proteins were stained for nuclei, IFITM3 (HA), early endosomes (EEA1), late endosomes (Rab7), and lysosomes (LAMP-1). Representative confocal images are shown. Arrowheads indicate regions of IFITM localization. Scale bars, 10 μm. For panels A–D, bars represent means from three independent experiments, with individual data points shown. Error bars indicate standard deviations. Statistical significance was determined by one-way ANOVA with multiple-comparison correction. *, P < 0.05; **, P < 0.01; ***, P < 0.001; ****, P < 0.0001.

Despite increased expression, disruption of the PPxY motif produced no discernible effects on HIV-1 infectivity (Fig. 5D), suggesting that enhanced protein levels alone is insufficient to substantially increase HIV-1 restriction. Confocal microscopy revealed that IFITM3-R1 AANY localized predominantly within late endosomal compartments (Fig. 5F), whereas both IFITM3-R2 and IFITM3-R2 AANY maintained diffuse distributions (Figs. 2L and 5G). Together, these results indicate that the PPxY motif constrains the antiviral potential of IFITM retrocopies by limiting protein accumulation and, in the case of IFITM3-R1, restricting endosomal localization.

### Lysine 51 contributes to the attenuated antiviral activity of IFITM3-R1

Sequence comparison of macaque IFITMs identified a lysine residue at position 51 that was unique to IFITM3-R1 (Fig. 6A). Given the central role of lysine residues in ubiquitin-mediated regulation (32), we investigated whether K51 contributes to the reduced expression and antiviral activity of IFITM3-R1.

**Figure 6.**
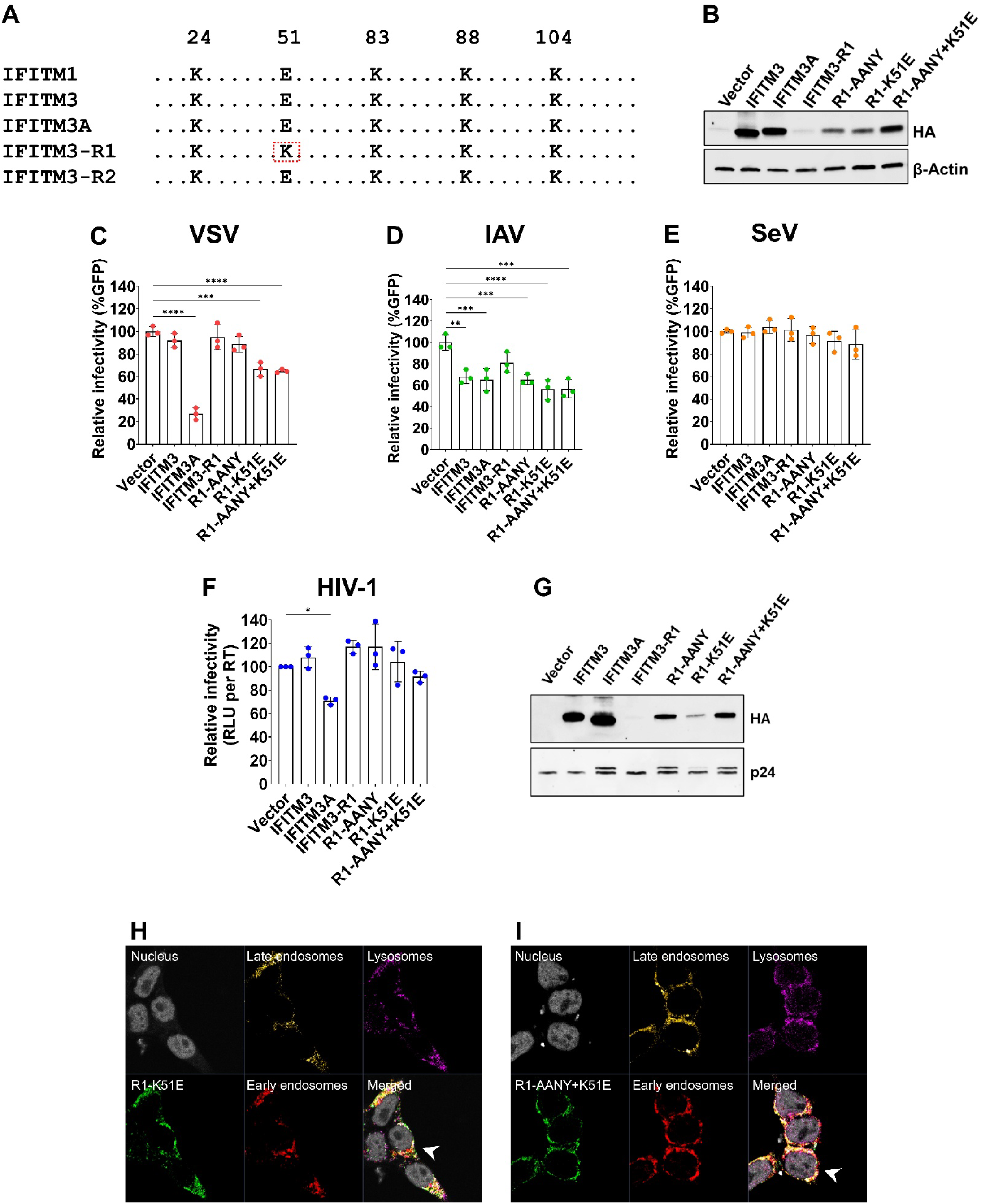
Residue K51 contributes to the reduced antiviral activity of IFITM3-R1. **(A)** Alignment of conserved lysine residues among canonical and non-canonical macaque IFITM proteins. The lysine residue unique to IFITM3-R1 is highlighted. **(B)** Mutation of the PPxY motif and K51E substitution increase IFITM3-R1 protein expression. HEK293T cells expressing HA-tagged IFITM3, IFITM3A, IFITM3-R1, or the indicated IFITM3-R1 mutants were analyzed by Western blot using anti-HA and anti-β-actin antibodies. **(C–E)** Effect of PPxY and K51E substitutions on antiviral activity of IFITM3-R1. HEK293T cells expressing the indicated IFITM proteins or mutants were infected with GFP-expressing VSV **(C)**, IAV **(D)**, or SeV **(E)**. Infection was quantified by flow cytometry as the percentage of GFP-positive cells and normalized to the empty vector control. **(F)** Effect of PPxY and K51E substitutions on HIV-1 infectivity. HIV-1 virions were produced in HEK293T cells expressing the indicated IFITM proteins or mutants. Infectivity was determined on TZM-bl target cells by measuring RLU normalized to RT activity and expressed relative to the empty vector control. **(G)** Incorporation of IFITM3-R1 mutants into HIV-1 virions. Virions produced in the presence of the indicated IFITM proteins were analyzed by Western blot using anti-HA and anti-p24 antibodies. **(H–I)** Subcellular localization of IFITM3-R1-K51E **(H)** and IFITM3-R1-AANY+K51E **(I)**. HEK293T cells expressing the indicated proteins were stained for nuclei, IFITM3 (HA), early endosomes (EEA1), late endosomes (Rab7), and lysosomes (LAMP-1). Representative confocal images are shown. Arrowheads indicate regions of IFITM localization. Scale bars, 10 μm. For panels C–F, bars represent means from three independent experiments, with individual data points shown. Error bars indicate standard deviations. Statistical significance was determined by one-way ANOVA with multiple-comparison correction. *, P < 0.05; **, P < 0.01; ***, P < 0.001; ****, P < 0.0001.

Substitution of lysine 51 with glutamic acid increased IFITM3-R1 expression to levels comparable to those observed following disruption of the PPxY motif (Fig. 6B). Combining the K51E substitution with the AANY mutation further enhanced protein levels, restoring expression to levels similar to those observed for canonical IFITM3 proteins.

Functional analysis revealed that K51E significantly enhanced restriction of both VSV and IAV infection (Fig. 6C–D). Notably, the antiviral activity of the K51E mutant against IAV was comparable to that observed with the PPxY mutant, indicating that either mutation alone is sufficient to enhance restriction of this virus. In contrast, K51E did not significantly affect SeV or HIV-1 infection (Fig. 6E–F), demonstrating that the effects of this residue are virus specific. Although the double mutant exhibited the highest protein expression, its antiviral activity was not uniformly greater than that of the individual mutants, suggesting that factors beyond protein abundance contribute to antiviral function.

Confocal microscopy revealed diffuse localization of both K51E and AANY+K51E mutants (Fig. 6H–I), indicating that enhanced antiviral activity does not require pronounced endosomal enrichment. Collectively, these findings identify K51 as a second determinant that contributes to the attenuated antiviral activity of IFITM3-R1.

### Differential incorporation of macaque IFITMs into HIV-1 virions

Because expression of both human and macaque IFITMs in target cells had little effect on HIV-1 infection (Figs. 1E and 2E), whereas expression in producer cells reduced virion infectivity (Figs. 1F and 2F), we next examined whether macaque IFITMs are incorporated into HIV-1 particles.

Consistent with previous reports (23–26), human IFITM1, IFITM2, and IFITM3 were readily detected in purified virions (Fig. 1G). Among macaque proteins, IFITM3 and IFITM3A exhibited the highest levels of incorporation, whereas IFITM1 and IFITM3-R2 were packaged less efficiently (Fig. 2G). IFITM3-R1 was not readily detected in virions under basal conditions, consistent with its low steady-state expression. However, disruption of the PPxY motif markedly enhanced incorporation of IFITM3-R1 into HIV-1 particles (Fig. 5E), and the combined AANY+K51E mutant exhibited greater virion incorporation than K51E alone (Fig. 6G).

Interestingly, increased incorporation did not always correlate with enhanced HIV-1 restriction. For example, PPxY disruption substantially increased incorporation of IFITM3-R1 into HIV-1 virions but had no observable effects on HIV-1 infectivity (Fig. 5D). Similarly, K51E enhanced IFITM3-R1 expression and packaging without significantly improving HIV-1 restriction (Fig. 6F). In contrast, IFITM3-R2 and IFITM3-R2 AANY had substantially lower virion incorporation but still displayed HIV-1 restriction comparable to IFITM3A (Figs. 5D–E). These findings suggest that incorporation into virions is necessary but not solely sufficient for antiviral activity and that distinct sequence determinants differentially regulate restriction of HIV-1 and other enveloped viruses.

Together, these results support a model in which macaque IFITMs inhibit HIV-1 primarily through incorporation into progeny virions rather than by restricting viral entry into target cells and demonstrate that evolutionary diversification of the macaque IFITM repertoire has generated proteins with distinct antiviral activities, intracellular trafficking patterns, and mechanisms of regulation.

## Discussion

IFITM proteins constitute a highly conserved family of interferon-stimulated restriction factors that inhibit the entry of a diverse range of enveloped viruses. While most studies have focused on the antiviral activities of human IFITM1, IFITM2, and IFITM3, comparative genomic analyses have revealed substantial lineage-specific diversification of IFITM genes across vertebrates (15). In particular, non-human primates exhibit striking differences in IFITM gene content, with several species encoding lineage-specific duplications, retrocopies, and pseudogenes (15–17). In our previous evolutionary analysis, we demonstrated that macaques possess a substantially expanded IFITM repertoire compared with humans, including a duplicated IFITM3 paralog (IFITM3A) and multiple retrotransposed IFITM3-derived genes (16). Here, we show that this expansion has generated proteins with distinct antiviral activities, intracellular trafficking patterns, and mechanisms of protein turnover, illustrating how gene duplication and retrotransposition can diversify innate antiviral defenses.

Expansion and diversification of restriction factor families is a recurring theme in host-virus evolution. Similar patterns have been described for APOBEC3 and Mx GTPases, where gene duplication events generated proteins with altered antiviral specificities or mechanisms of action. The IFITM family appears to follow a similar evolutionary trajectory. Although humans encode three antiviral IFITM proteins, many mammalian species contain expanded IFITM loci. Rodents and marsupials possess additional IFITM paralogs (15, 33), bats exhibit extensive IFITM diversification driven by gene duplication and alternative splicing that is associated with antiviral adaptation (34, 35), and several non-human primates encode lineage-specific IFITM duplications (15–17). The presence of both duplicated and retrotransposed IFITM3 genes in macaques suggests recurrent selective pressure to expand IFITM-mediated antiviral defenses. Our findings indicate that these duplicated genes have not simply been retained as redundant copies but instead have acquired distinct functional properties.

In this study, among the macaque IFITMs examined, IFITM3A consistently displayed the strongest antiviral activity, particularly against VSV. Although IFITM3A originated from duplication of IFITM3, the two proteins differ at multiple amino acid positions and exhibit distinct intracellular localization patterns. Whereas IFITM3 localized predominantly to Rab7-positive late endosomes, IFITM3A displayed a broader distribution encompassing both early and late endosomal compartments. These observations are notable because IFITM antiviral activity is influenced by subcellular localization. Previous studies demonstrated that IFITM1 primarily restricts viruses that fuse at or near the plasma membrane, whereas IFITM2 and IFITM3 preferentially inhibit viruses that enter through endosomal pathways (4, 5, 17, 36–40). The broader endosomal distribution of IFITM3A may therefore increase its ability to intercept incoming viral particles across multiple stages of the endocytic pathway and could explain its enhanced activity against VSV.

To investigate the molecular basis for this gain of function, we selected amino acid residues that differed between macaque IFITM3A and IFITM3 and that occurred within regions previously implicated in IFITM trafficking and antiviral activity. Sequence comparisons among macaque IFITM3A, macaque IFITM3, and human IFITM2 identified three candidate substitutions: F8V within the N-terminal domain, P70T within the intramembrane domain, and I108V within the C-terminal region. Notably, both F8V and P70T convert IFITM3 residues to amino acids shared by IFITM3A and human IFITM2, suggesting that these substitutions may represent convergent determinants of altered antiviral activity or intracellular trafficking. In contrast, I108V was selected because previous studies implicated the neighboring 103–108 region in regulating IFITM stability and incorporation into HIV-1 virions (29). Functional analyses revealed that F8V and P70T preferentially enhanced restriction of VSV, whereas I108V selectively increased anti-HIV activity. These findings indicate that distinct regions of IFITM3 contribute to restriction of different viruses and suggest that the enhanced antiviral phenotype of IFITM3A arose through the combined effects of multiple amino acid substitutions rather than a single gain-of-function mutation. More broadly, these results illustrate how relatively small sequence changes following gene duplication can diversify antiviral specificity while preserving the overall restriction factor architecture. Similar virus-specific effects have been reported for human IFITM3 polymorphisms, including variants that alter susceptibility to influenza virus and SARS-CoV-2 infection (41). Together, these observations support a model in which IFITM proteins evolve through incremental sequence changes that fine-tune antiviral specificity against distinct viral pathogens.

Another key finding of this study is that retrotransposed IFITM genes remain functionally active despite substantial divergence from canonical IFITM proteins. Retrogenes are frequently viewed as evolutionary dead ends because they lack regulatory elements and often accumulate inactivating mutations (42). Nevertheless, several antiviral retrogenes have been shown to acquire novel functions following retrotransposition, including retrocopies of APOBEC3 (43), TRIMCyp fusion (44), and RetroCHMP3 (45). We found that both IFITM3-R1 and IFITM3-R2 retain antiviral activity, although their activities differ from those of canonical IFITM3 proteins. These results indicate that retrotransposition can contribute to expansion of the IFITM antiviral repertoire and suggest that non-canonical IFITM genes may serve as a reservoir for evolutionary innovation.

Our study also identifies a previously unrecognized mechanism that regulates IFITM retrocopy expression. IFITM3-R1 exhibited markedly reduced steady-state protein levels, which were restored by disruption of an N-terminal PPxY motif and by treatment with the lysosomal inhibitor chloroquine. PPxY motifs are well-established binding sites for WW-domain-containing proteins, including members of the NEDD4 family of E3 ubiquitin ligases (30, 31). Human IFITM3 is regulated by NEDD4-mediated ubiquitination, which promotes protein turnover and limits antiviral activity (46). Our findings suggest that a similar pathway regulates IFITM3-R1, but with substantially greater impact on protein stability. The observation that chloroquine, but not MG132, restored IFITM3-R1 expression further indicates that lysosomal degradation represents the dominant pathway controlling turnover of this retrocopy. Thus, diversification of IFITM genes appears to involve not only changes in antiviral function but also evolution of distinct mechanisms governing protein stability.

Interestingly, increased IFITM expression did not uniformly translate into enhanced antiviral activity. Mutation of the PPxY motif dramatically increased IFITM3-R1 expression and virion incorporation but produced little effect on HIV-1 restriction. Likewise, introduction of the K51E substitution increased expression and virion incorporation without measurably enhancing HIV-1 restriction. These observations suggest that IFITM abundance alone does not determine antiviral activity. Instead, specific sequence determinants likely influence how IFITMs alter membrane properties, interact with viral glycoproteins, and/or modulate membrane fusion events. Similar dissociation between IFITM expression levels and antiviral activity has been reported for human IFITM proteins (47), indicating that antiviral function depends on both abundance and intrinsic biochemical properties.

One of the most notable findings of this study is the striking difference between HIV-1 restriction in target cells and producer cells. Neither human nor macaque IFITMs inhibited HIV-1 infection when expressed in target cells, despite effectively restricting VSV and IAV. In contrast, several IFITMs reduced HIV-1 infectivity when expressed in virus-producing cells. These observations are consistent with a growing body of literature indicating that HIV-1 is relatively resistant to IFITM-mediated inhibition during target-cell entry but remains susceptible to IFITM-dependent alterations of virion infectivity. Multiple studies have demonstrated that IFITMs can be incorporated into budding HIV-1 particles, where they impair viral fusion and reduce infectivity of progeny virions (23–26). IFITMs have also been reported to alter Env processing, Env incorporation, and membrane fusogenicity, thereby reducing viral spread (26).

Our virion incorporation experiments support this producer-cell model but also reveal important complexities. IFITM3 and IFITM3A were incorporated efficiently into HIV-1 particles, whereas IFITM3-R2 displayed comparatively lower incorporation. Surprisingly, virion incorporation efficiency did not correlate directly with antiviral activity. For example, disruption of the PPxY motif greatly enhanced incorporation of IFITM3-R1 into virions without significantly increasing HIV-1 restriction. Conversely, IFITM3-R2 restricted HIV-1 despite relatively low levels of virion incorporation. These findings suggest that IFITM virion incorporation is necessary but not sufficient for anti-HIV-1 activity and that qualitative features of incorporated IFITMs may be more important than absolute abundance. Previous studies similarly reported that virion-associated IFITM levels do not always predict the magnitude of HIV-1 inhibition and proposed that specific IFITM-Env interactions or alterations in membrane architecture may determine antiviral activity (18, 29, 48, 49). Our data are consistent with this model and suggest that evolutionary diversification of macaque IFITMs has uncoupled virion incorporation from antiviral efficacy.

Finally, our localization studies provide additional insight into the relationship between intracellular trafficking and antiviral specificity. IFITM proteins that localized predominantly to endosomal compartments generally exhibited the strongest activity against viruses that utilize endocytic entry pathways, such as VSV and IAV. In contrast, IFITM3-R2 displayed a more diffuse distribution and comparatively weak antiviral activity against these viruses. However, localization alone could not fully explain antiviral phenotypes. For example, K51E mutants retained diffuse localization while exhibiting enhanced antiviral activity. Thus, although endosomal localization likely contributes to antiviral potency, additional determinants such as membrane composition, protein turnover, oligomerization, and/or interactions with host trafficking machinery may influence restriction activity.

In summary, our findings demonstrate that expansion of the macaque IFITM locus has generated a functionally diverse antiviral repertoire through both gene duplication and retrotransposition. The duplicated paralog IFITM3A acquired enhanced antiviral activity accompanied by altered intracellular localization, whereas the retrocopies IFITM3-R1 and IFITM3-R2 evolved distinct regulatory and antiviral properties. We further identify the PPxY motif and residue K51 as determinants that limit expression and antiviral activity of IFITM3-R1 and show that macaque IFITMs modulate HIV-1 infectivity predominantly through producer-cell-dependent mechanisms. Although several macaque IFITMs are efficiently incorporated into HIV-1 virions, virion incorporation alone does not predict antiviral activity, indicating that additional cis- or trans-acting determinants contribute to HIV-1 restriction. Together, these findings provide a mechanistic framework for understanding how IFITM diversification expands innate antiviral defenses and illustrate how evolutionary innovation within restriction factor families generates proteins with distinct antiviral functions and regulatory mechanisms.

## Materials and Methods

### Cells, plasmids, viruses

HEK293T (ATCC CRL-3216), HEK293T expressing human CD4 and CCR5 (50), HeLa TZM-bl (51) (BEI Resources, NIH HIV Reagent Program, catalog no. 8129), MDCK (ATCC CCL-34), and Vero (ATCC CCL-81) cells were cultured in Dulbecco’s modified eagle medium (DMEM, Gibco) supplemented with 10% fetal bovine serum (FBS, Sigma), 2 mM L-glutamine (Gibco), and 1x Penicillin-Streptomycin (Gibco) (complete DMEM).

The following plasmids encoding N-terminally HA-tagged human IFITM proteins were used: pCMV-HA-hIFITM1 (Addgene plasmid no. 58399), pCMV-HA-hIFITM2 (Addgene plasmid no. 58398), and pCMV-HA-hIFITM3 (Addgene plasmid no. 58397). Open reading frames encoding N-terminally HA-tagged macaque IFITM1, IFITM3, IFITM3A, IFITM3-R1, and IFITM3-R2 were synthesized as DNA fragments (Twist Bioscience), digested, and ligated into pCMV-HA (Takara Bio) using the EcoRI and SalI restriction sites to generate pCMV-HA macaque IFITM variant expression plasmids. Plasmids encoding macaque IFITM3 mutants were generated using the QuikChange Multi Site-Directed Mutagenesis Kit (Agilent). All plasmids generated in this study were verified by Sanger DNA sequencing.

Green fluorescent protein (GFP)-reporter HIV-1 pseudoviruses were generated as described previously (52). Briefly, HEK293T cells were cotransfected with 4 µg of Envelope (Env)-deficient HIV-1 proviral plasmid (Q23ΔEnvGFP) and 2 µg of HIV-1_BG505_ or HIV-1_NL4-3_ Env expression plasmids using Fugene 6 transfection reagent (Roche) following manufacturer’s protocol. Pseudovirus titers were determined by infection of TZM-bl cells, followed by staining for β-galactosidase activity at 48 hours post-infection as described previously (51). Influenza virus A/Puerto Rico/8/1934 (H1N1, PR8) expressing GFP (rIAV-GFP) was provided by Dr. Adolfo Garcia-Sastre (Icahn School of Medicine at Mt. Sinai) and was propagated in 10-day embryonated chicken eggs for 48 hours at 37°C and titrated in MDCK cells as described previously (53). Recombinant vesicular stomatitis virus (VSV) Indiana strain expressing GFP (rVSV-GFP) and Sendai virus expressing GFP (rSeV-GFP) were propagated and titrated in Vero cells by plaque assay and TCID_50_ assay, respectively, as described previously (54).

### Infections of IFITM-expressing target cells

HEK293T cells were seeded at a density of 2.5 x 10^5^ cells per well in 6-well plates 24 hours prior to transfection. Cells were transfected with plasmids encoding human or macaque HA-tagged IFITM variants, or with an empty vector control, using Fugene 6 transfection reagent (Roche) following manufacturer’s protocol. At 24 hours post-transfection, cells were infected with rIAV-GFP at a multiplicity of infection (MOI) of 1.25 for 8 hours, rVSV-GFP at an MOI of 0.005 for 12 hours, or rSeV-GFP at an MOI of 0.5 for 12 hours. For rSeV-GFP infections, FBS-containing complete DMEM was replaced with fresh DMEM supplemented with 0.5 µg/mL TPCK-treated trypsin prior to infection. Following infection, cells were harvested using 0.05% trypsin-EDTA (Gibco), washed twice with 1x phosphate-buffered saline (PBS), fixed with 4% paraformaldehyde in 1x PBS for 10 minutes, and permeabilized with 0.1% Triton X-100 in 1x PBS for 10 minutes. Cells were then stained with anti-HA.11 primary antibody (1:250, BioLegend catalog no. 901521) diluted in fluorescence-activated cell sorter (FACS) buffer (1x PBS, 1% FBS, 1 mM EDTA) for 20 minutes at room temperature. After two washes with FACS buffer, cells were incubated for 20 minutes at room temperature in the dark with Alexa Fluor 647-conjugated goat anti-mouse secondary antibody (1:500, Invitrogen catalog no. A21235) diluted in FACS buffer. Cells were washed twice with FACS buffer, resuspended in FACS buffer, and analyzed for GFP and Alexa Fluor 647 expression using an Attune NxT flow cytometer. Data were analyzed using FlowJo software.

For HIV-1 infection of IFITM-expressing target cells, the above procedure was adapted using HEK293T cells expressing human CD4 and CCR5 receptors. Briefly, 1.25 x 10^5^ cells were seeded per well in 6-well plates and transfected with plasmids encoding human or macaque HA-tagged IFITM variants, or with an empty vector control, using Fugene 6 transfection reagent (Roche) following manufacturer’s protocol. At 24 hours post-transfection, cells were infected with GFP-reporter HIV-1 pseudoviruses at an MOI of 1 in the presence of 10 μg/ml of DEAE-dextran for 48 hours. Following infection, cells were processed for anti-HA staining and GFP expression analysis by flow cytometry as described above.

### Infectivity of HIV-1 pseudoviruses produced from IFITM-expressing cells

HEK293T cells were seeded at a density of 2.5 x 10^5^ cells per well in 6-well plates 24 hours prior to transfection. Cells were cotransfected with plasmids encoding Env-deficient HIV-1 proviral backbone, HIV-1 Env, and human or macaque HA-tagged IFITM variants, or an empty vector control, using Fugene 6 transfection reagent (Roche) following manufacturer’s protocol. At 24 hours post-transfection, culture supernatants were replaced with 2 ml of fresh complete DMEM. After an additional 24 hours, pseudovirus-containing supernatants were clarified by centrifugation at 650 x g for five minutes at room temperature, passed through a 0.2 μm sterile filter, and aliquoted for downstream analyses.

HIV-1 pseudovirus infectivity was measured using TZM-bl indicator cells and the Galacto-Light Plus β-Galactosidase Assay (ThermoFisher), following manufacturer’s protocol, and normalized to reverse transcriptase (RT) activity. Briefly, TZM-bl cells were seeded at a density of 1 x 10^4^ cells per well in black 96-well polystyrene plates in 100 µL of complete DMEM and infected with HIV-1 pseudoviruses in the presence of 15 μg/ml of DEAE-dextran. At 48 hours post-infection, supernatants were removed, cells were washed once with 1x PBS, and lysed in 10 µL of lysis solution for 10 minutes at room temperature. Subsequently, 70 µL per well of reaction buffer containing a 1:100 dilution of Galacton-Plus substrate was added and incubated for 50 minutes in the dark. Next, 100 µL of accelerator solution was added, and luminescence was measured using an Infinite M Plex multimode microplate reader.

### Reverse transcriptase activity assay

RT activity assay was performed as described previously (55). Briefly, 5 µl of HIV-1 pseudoviral stock or RT standard was lysed in 5 µl of 2x lysis buffer (100 mM Tris HCl pH 7.4, 50 mM KCl, 0.25% Triton X-100, 40% glycerol) in the presence of 4U RNaseOUT (Invitrogen) for 10 minutes at room temperature. Viral lysate was diluted 1:10 by adding 90 µl of nuclease-free water (Life Technologies). qRT-PCR reactions were prepared by mixing 9.6 µl of diluted viral lysate with 10.4 µl of reaction mix containing 10 µl of 2x Maxima SYBR Green/ROX qPCR Master Mix (ThermoFisher), 0.1 µl of 4U/µl RNaseOUT, 0.1 µl of 0.8 µg/µl MS2 RNA template (Roche), and 0.1 µl each of 100 µM forward 5’-TCCTGCTCAACTTCCTGTCGAG-3’ and reverse 5’-CACAGGTCAAACCTCCTAGGAATG-3’ primers. qRT-PCR was performed using a QuantStudio 3 Real-Time PCR machine (Applied Biosystems). Viral titers were calculated from a standard curve generated using recombinant reverse transcriptase (Millipore catalog no. 382129).

### Immunoblotting

Whole cell extracts were prepared by lysing the cells in radioimmunoprecipitation assay (RIPA) cell lysis buffer (50 mM Tris pH 8.0, 0.1% SDS, 1% Triton-X, 150 mM NaCl, 1% deoxycholic acid, 2 mM PMSF). For IFITM incorporation in HIV-1 virions, virus containing supernatants from the infected cell cultures were centrifuged at 650 x g for five minutes at room temperature. Cell-free supernatant was filtered through 0.2 µm filter and then pelleted through a 25% sucrose cushion by ultracentrifugation for at 28,000 rpm for 90 minutes at 4°C. Virus pellets were lysed in 70 μl of RIPA buffer for 10 minutes at room temperature. The concentration of HIV-1 Gag in the viral lysates was determined by RT assay, and normalized amounts of lysate were subjected to SDS-PAGE and immunoblotted. Standard Western blotting procedures were used with the following antibodies: HA.11 epitope tag (BioLegend catalog no. 901521), HIV-1 p24 (BEI Resources, NIH HIV Reagent Program catalog no. 6521), and β-Actin (Abcam catalog no. ab3280).

### Protein degradation assay

HEK293T cells were seeded at a density of 2.5 x 10^5^ cells per well in 6-well plates 24 hours prior to transfection. Cells were transfected with plasmids encoding macaque IFITM3 variants or an empty vector control using Fugene 6 transfection reagent (Roche) following manufacturer’s protocol. At 2 hours post-transfection, cells were treated with either the proteasomal inhibitor MG132 (10 μM) or the lysosomal inhibitor chloroquine (10 μM) for 24 hours. Following treatment, cells were washed with 1x phosphate-buffered saline (PBS) and harvested using 0.05% trypsin-EDTA (Gibco). Cells were subsequently washed again with 1x PBS and whole cell extracts were prepared using RIPA lysis buffer. Protein lysates were then analyzed by immunoblotting.

### Immunofluorescent Microscopy

For imaging experiments, 10^5^ HEK293T cells were cultured on sterilized glass coverslips in 12-well plates and transfected with plasmids encoding human or macaque HA-tagged IFITM variants, or an empty vector control, using Fugene 6 transfection reagent (Roche) following manufacturer’s protocol. At 24 hours post-transfection, cells were fixed with 4% paraformaldehyde in 1x phosphate-buffered saline (PBS) for 15 minutes and permeabilized with 0.1% Triton X-100 in 1x PBS containing 2% bovine serum albumin (BSA) for 10 minutes. Cells were incubated overnight at 4°C in a humidified chamber with anti-HA.11 primary antibody (1:1000, BioLegend catalog no. 901521). Following two washes with 1x PBS, cells were incubated for 6 hours at 4°C in a humidified chamber with Alexa Fluor 647-conjugated goat anti-mouse secondary antibody (1:500, Invitrogen catalog no. A21235) together with antibodies against early endosomes (Alexa Fluor 594 Anti-EEA1, 1:1000, Abcam catalog no. 206913), late endosomes (Alexa Fluor 488 Anti-RAB7, 1:500, Abcam catalog no. 309763), or lysosomes (eFluor 450 Anti-LAMP-1, 1:400, ThermoFisher catalog no. 48107942 ), as indicated. Cells were subsequently washed twice with 1x PBS and stained with Chromomycin A3 (ThermoFisher) for 30 minutes at 4°C in a humidified chamber. Coverslips were washed twice with 1x PBS, mounted in ProLong Diamond Antifade reagent (Invitrogen), and cured overnight at room temperature in the dark. Images were acquired using a Zeiss LSM 880 confocal microscope and analyzed with ZEN Black software.

**Supplementary Figure S1. Genomic organization of canonical and non-canonical IFITM loci in rhesus macaques and humans. (A)** Syntenic comparison of the canonical IFITM locus in macaques (chromosome 14) and humans (chromosome 11), showing the arrangement of IFITM family members and neighboring genes. **(B)** Genomic location of the macaque IFITM3-R1 retrocopy on chromosome 17 relative to the syntenic human locus. **(C)** Genomic location of the macaque IFITM3-R2 retrocopy on chromosome 9 relative to the syntenic human locus. Colored boxes indicate canonical IFITM genes, IFITM retrocopies, and exons derived from the parental PFAS and MCU genes. Black arrows indicate gene orientation and transcriptional direction.

**Supplementary Figure S2. Alignment of human and macaque IFITM proteins.** Amino acid sequence alignment of canonical and non-canonical IFITM proteins from humans and macaques. Human IFITM1, IFITM2, and IFITM3 were aligned with macaque IFITM1, IFITM3, IFITM3A, IFITM3-R1, and IFITM3-R2 using Clustal Omega. Conserved residues are indicated in the consensus sequence shown at the top. Dashes represent alignment gaps introduced to maximize sequence similarity. The alignment highlights the overall conservation of canonical IFITM proteins across species, as well as sequence divergence within the macaque-specific IFITM3A paralog and the non-canonical retrocopies IFITM3-R1 and IFITM3-R2. Notably, IFITM3-R2 encodes a truncated protein lacking the conserved C-terminal transmembrane domain present in canonical IFITM proteins.

**Supplementary Figure S3. Sequence conservation and divergence among macaque IFITM3 proteins associated with altered antiviral activity.** Sequence logo analysis of macaque IFITM3A, macaque IFITM3, human IFITM3, and human IFITM2. Amino acid conservation at each position is represented by letter height, with the total stack height indicating the degree of conservation. Residues that vary among the four proteins are shown below the sequence logo. The analysis highlights a limited number of amino acid differences distributed throughout the proteins, including substitutions previously implicated in regulating IFITM trafficking, membrane localization, and antiviral activity. The high overall conservation between macaque IFITM3A and IFITM3 indicates that functional differences between these paralogs arise from a small number of sequence changes rather than extensive divergence across the protein.

## Acknowledgments

We thank Julie Overbaugh for providing HEK293T expressing human CD4 and CCR5 and Adolfo Garcia-Sastre for providing rIAV-GFP. The following reagents were obtained through BEI Resources, NIAID, NIH: TZM-bl cells and anti-HIV-1 p24 Gag monoclonal antibody.

